# Profiling Epigenetic Aging at Cell-Type Resolution Through Long-Read Sequencing

**DOI:** 10.1101/2024.11.20.623937

**Authors:** Alec Eames, Mahdi Moqri, Jesse R. Poganik, Vadim N. Gladyshev

## Abstract

DNA methylation can give rise to robust biomarkers of aging, yet most studies profile it at the bulk tissue level, which masks cell type-specific alterations that may follow distinct aging trajectories. Long-read sequencing technology enables methylation profiling of extended DNA fragments, which allows mapping to their cell type of origin. In this study, we introduce a framework for evaluating cell type-specific aging using long-read sequencing data, without the need for cell sorting. Leveraging cell type-specific methylation patterns, we map long-read fragments to individual cell types and generate cell type-specific methylation profiles, which are used as input to a newly developed probabilistic aging model, LongReadAge, capable of predicting epigenetic age at the cell-type level. We apply LongReadAge to track aging of myeloid cells and lymphocytes from bulk leukocyte data as well as circulating cell-free DNA, demonstrating robust performance in predicting age despite limited shared features across samples. This approach provides a novel method for profiling the dynamics of epigenetic aging at cell-type resolution.

## Introduction

Aging is characterized by progressive physiological decline and increased incidence of disease, disability, and death^1^. At the molecular level, epigenetic modifications have emerged as a key biomarker of aging, with widespread shifts observed across the epigenomic landscape as we age^2,3^. The most widely used epigenetic modification in aging research is methylation of cytosines at cytosine-guanine (CpG) dinucleotides, whose patterns enable robust prediction of age, mortality risk, and disease susceptibility^4–8^.

Emerging evidence suggests that aging involves cell type-specific alterations ^9–13^, underscoring the importance of tracking aging at cell-type resolution. Existing methylation-based biomarkers of aging, however, largely rely on bulk methylation profiles, rendering them susceptible to confounding from cell composition and obscuring signals from low-abundance cell types ^14^. While algorithms to computationally generate cell-type methylation patterns from bulk profiles have been developed^15^, these approaches are hindered by their dependence on known cell proportions. Cell-type methylation patterns can be determined experimentally through cell sorting and single-cell sequencing; however, such techniques are time-consuming, expensive, and not scalable for routine use.

With the advent of long-read sequencing technology, it is now possible to sequence extended fragments of DNA in a single read^16^. Crucially, this technology enables the simultaneous determination of nucleotide sequence and CpG methylation, without the need for experimental cytosine-to-uracil conversion. Excitingly, recent studies have demonstrated the feasibility of computationally tracing fragments from long-read sequencing to their cell type of origin using cell type-specific methylation patterns^17–19^, obviating the need for cell sorting. A drawback of these long-read methylation profiles, however, is the small number of intersecting CpG sites across samples, which impedes across-sample age profiling, particularly for cell type-specific methylation profiles derived from a limited number of reads.

In this study, we leverage long-read methylation data paired with a newly developed probabilistic aging model, LongReadAge, to profile epigenetic aging at cell-type resolution. LongReadAge, based on the scAge framework^13^, computes the epigenetic age of sparsely intersecting long-read methylation profiles. By mapping fragments from long-read data to their cell type of origin and applying LongReadAge to the resulting methylation profiles, we show that cell type-specific estimates of epigenetic age may be produced from long-read sequencing of bulk samples. We focus our analysis on peripheral blood, a widely studied tissue composed of a heterogeneous mixture of cell types, through which we track aging of myeloid cells and lymphocytes––two fundamental hematopoietic cell types that display distinct features during aging^20,21^. We further apply our model to track aging of circulating cell-free DNA (Figure 1).

**Figure 1:**
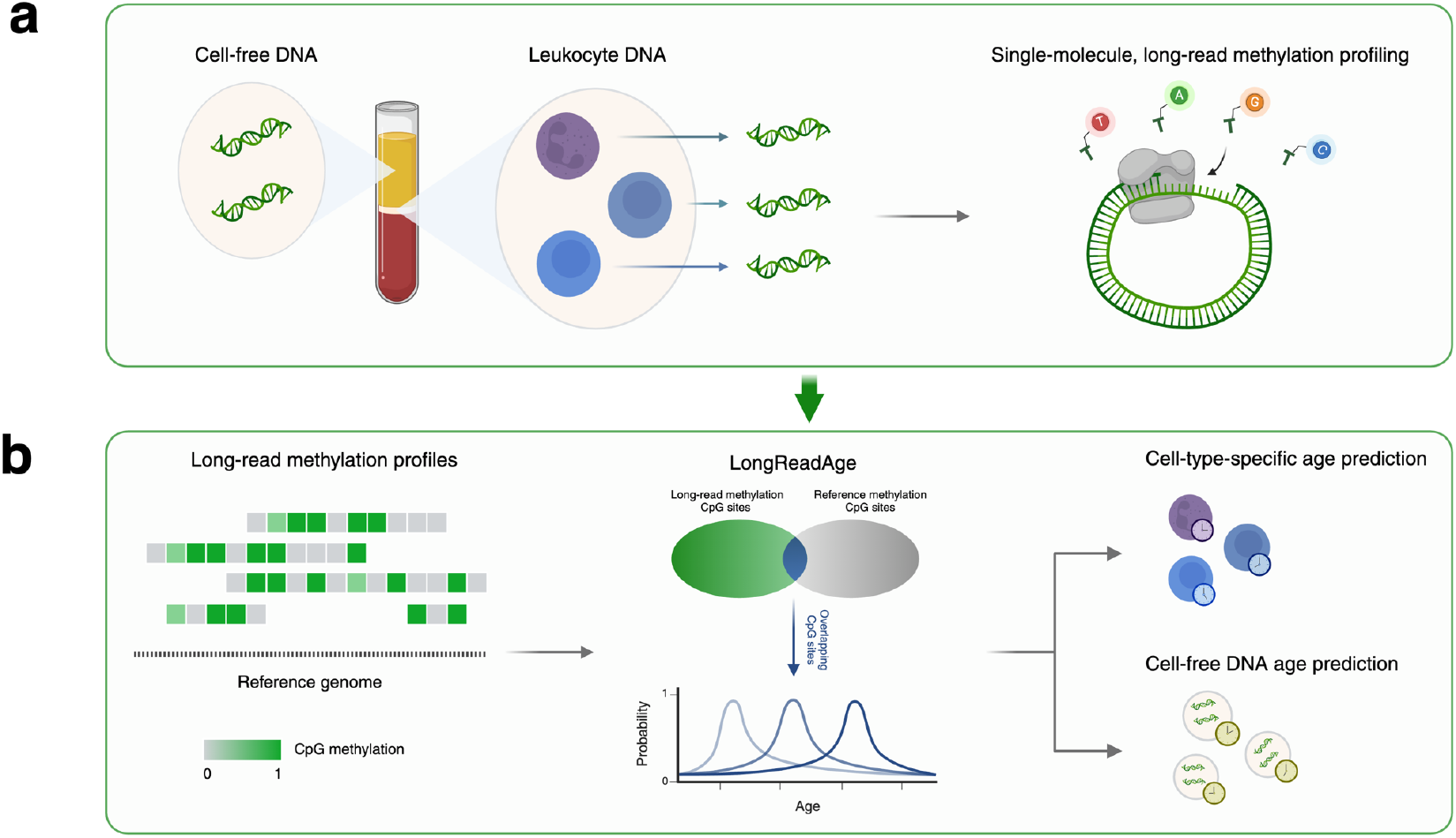
Schematic overview of the study. **(a)** CpG methylation of peripheral leukocytes and circulating cell-free DNA was analyzed through long-read sequencing. **(b)** A probabilistic model, LongReadAge, was developed to predict epigenetic age using sparsely intersecting long-read methylation profiles. We applied this model to profile immune-cell-specific and cell-free DNA aging.

## Results

### Development and application of LongReadAge

Inspired by the scAge framework^13^, we developed LongReadAge, a probabilistic model designed to predict epigenetic age based on long-read methylation data (Figure 1). In this model, a reference aging dataset is used to build linear regression models predicting methylation as a function of age across every profiled CpG site. The input to LongReadAge is a long-read methylation profile, consisting of a series of CpG sites along with their methylation status. We identify CpGs common to both the input long-read profile and the reference dataset, then filter these CpGs to include only those with the highest Pearson correlation with chronological age in the reference dataset. Next, for this filtered set of CpGs, we predict methylation patterns over a range of ages, from 0 to 100 years, using the linear age-methylation relationships found in the reference dataset. For each age’s predicted methylation profile, we compare it to the input long-read methylation profile using a probability score that quantifies the similarity between the two profiles (see Methods). The ages that yield the highest probability score, indicating the closest match between the predicted and observed long-read profile, is considered the epigenetic age of the sample (Figure 2A).

**Figure 2:**
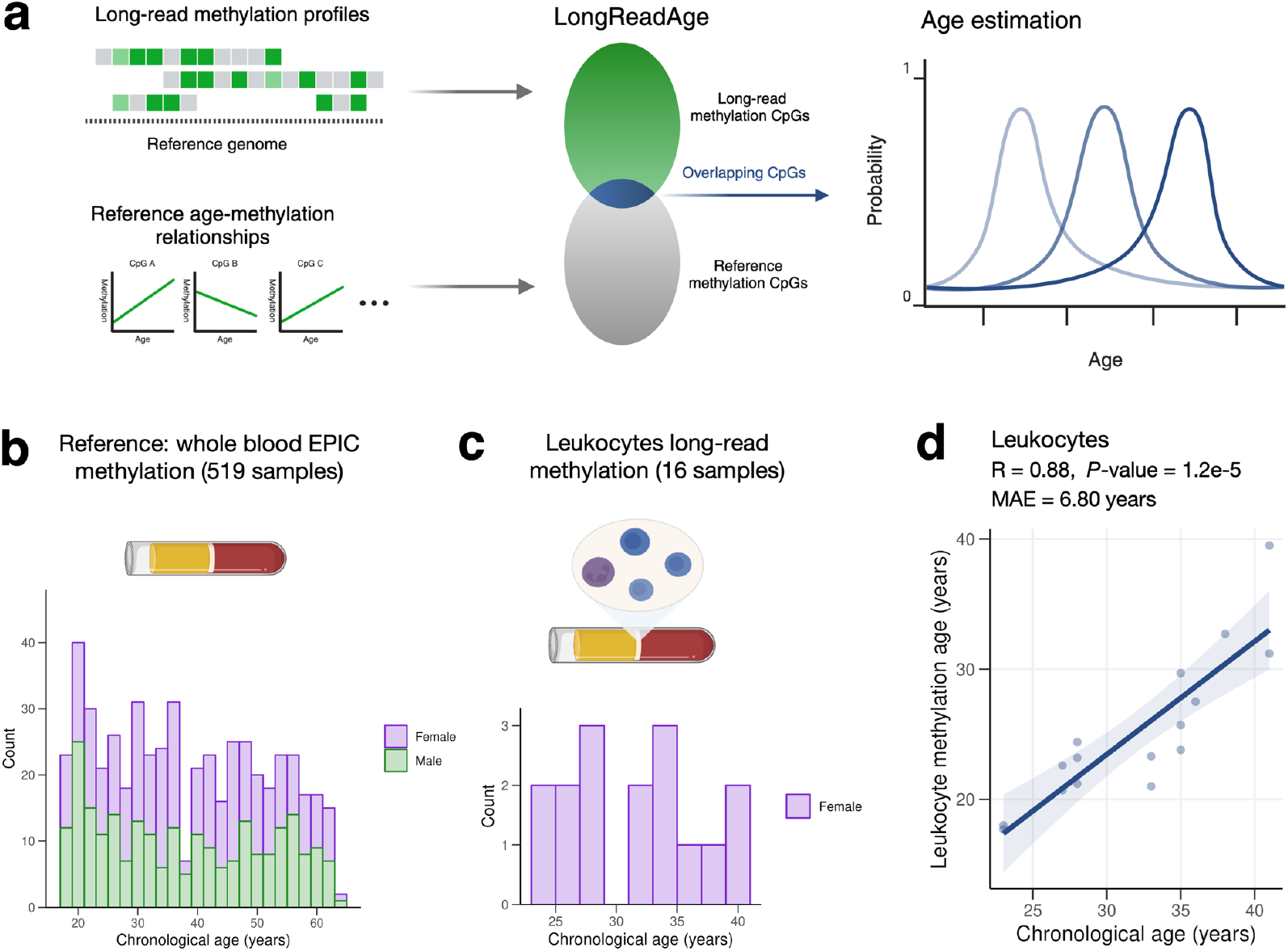
Development of LongReadAge. **(a)** Schematic depicting LongReadAge framework. In this model, CpG sites from long-read methylation profiles are intersected with CpGs from a reference dataset, where the relationship between chronological age and CpG methylation is well-characterized. For these intersecting CpG sites, methylation status is predicted using chronological age. The age that best predicts the observed long-read methylation pattern is considered the epigenetic age of the sample. **(b)** Our reference dataset used in this model consisted of CpG methylation from 519 healthy individuals analyzed using the EPIC microarray platform. **(c)** Long-read single-molecule methylation sequencing data was obtained from 16 samples. DNA was extracted from peripheral leukocytes (buffy coat). **(d)** Application of LongReadAge to long-read leukocyte data.

We evaluated our model by applying it to data from a cohort of 11 pregnant women for which 16 long-read methylation profiles of peripheral leukocytes were generated in a previous study (Figure 2C)^18^. As our reference aging dataset, we used a whole-blood methylation dataset comprising 519 healthy individuals balanced between males and females measured with the EPIC microarray^22^ (Figure 2B). LongReadAge yielded epigenetic age estimations that displayed a strong correlation with chronological age and a mean absolute error below 7 years, demonstrating the potential of our approach in estimating age from sparsely intersecting long-read methylation data (Figure 2D). Notably, since LongReadAge is trained on the reference dataset, it can be applied to long-read datasets with a small number of subjects, as is the case here.

### LongReadAge captures immune-cell-specific aging

We next applied LongReadAge to profile cell type-specific aging. Beginning with the same 16 long-read bulk samples measuring leukocyte methylation, we generated cell-type methylation profiles by tracing individual fragments to their cell type of origin, following a previously developed and validated approach^17,18^. For this, we leveraged a comprehensive methylation atlas measuring genome-wide methylation across a range of cell types^23^. From this atlas, we identified differentially methylated CpG sites distinguishing two distinct cell types found in blood: lymphocytes and myeloid cells. We identified these sites using the *wgbstools* suite^24^, applying several criteria to identify robust differentially methylated CpG sites.

Next, we overlapped each long-read fragment’s CpG sites with the differentially methylated CpG sites in the reference atlas. Among fragments with at least five overlapping reference CpG sites, we computed a score describing the likelihood of a fragment originating from each cell type based on how well the fragment’s methylation patterns matched each respective cell type. Fragments with a 1.05x greater likelihood of originating from one cell type versus another were considered to derive from that cell type.

We aggregated all fragments corresponding to a given cell type to generate cell type-specific methylation profiles (Figure 3A). The accuracy of generating cell type-specific methylation profiles was verified with two approaches. First, we confirmed the proportion of reads predicted to originate from myeloid cells and lymphocytes matched typical proportions of these cell types in peripheral blood (Figure 3B)^25^. Among prediction-eligible fragments, approximately 75% were predicted to originate from myeloid cells, and around 20% were predicted to originate from lymphocytes, with a slight underrepresentation due to a conservative 1.05x probability cutoff. Second, we compared long-read cell type-specific methylation to reference cell-type methylation profiles, which revealed a strong concordance between the two. With methylation represented as a value ranging from 0 to 1 (universally unmethylated to universally methylated), the long-read lymphocyte profile displayed a median deviation of approximately 0.1 from the reference lymphocyte profile versus about 0.45 from the reference myeloid profile. The long-read myeloid profile displayed similar results, matching the myeloid reference much more closely than the lymphocyte reference (Figure 3C). Visual examination of long-read aggregated methylation profiles versus cell type-specific reference profiles underscored their similarity (Figure 3D).

**Figure 3:**
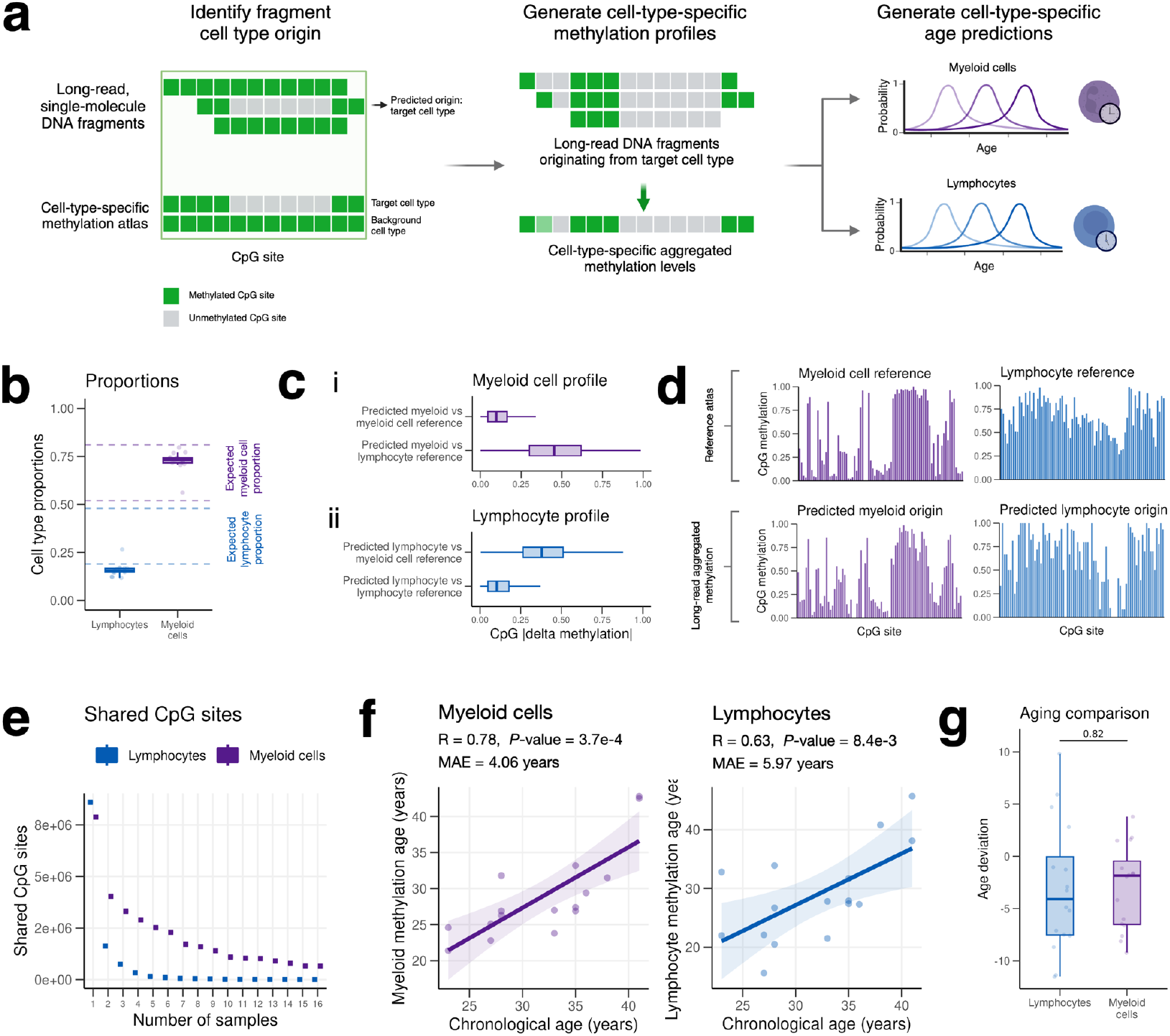
LongReadAge captures cell type-specific aging. **(a)** Schematic depicting cell type-specific methylation profile generation and age prediction. Using single-molecule long-read DNA methylation data from leukocytes, we identified the cellular origin of individual DNA fragments and obtained cell type-specific methylation profiles. These profiles were applied as input to LongReadAge to generate cell type-specific epigenetic age predictions. **(b)** Proportions of reads predicted to originate from lymphocytes and myeloid cells. Dotted lines depict ranges of typical lymphocyte and myeloid cell proportions in peripheral blood. **(c)** Average magnitude of CpG methylation difference between i) predicted myeloid cell methylation profile vs myeloid cell reference and vs lymphocyte reference, and ii) predicted lymphocyte methylation profile vs myeloid cell reference and vs lymphocyte reference. **(d)** Example methylation profiles from cell-specific methylation atlas (myeloid cells and lymphocytes) and averaged long-read methylation data for reads predicted to originate from myeloid and lymphocyte cells. Displayed here is a random selection of CpGs present in both lymphocyte and myeloid differentially methylated CpG sites and in at least one sample of myeloid- and lymphocyte-predicted long-read methylation profiles. **(e)** Number of shared CpGs between progressively increasing numbers of samples for both 1) lymphocyte long-read methylation profiles and 2) myeloid cell long-read methylation profiles. **(f)** Results of applying LongReadAge to cell type-specific methylation profiles. Two-step percentile, fixed number cutoff is used here to generate predictions. **(g)** Age deviation comparison between lymphocytes and myeloid cells for the same individuals.

Given that only reads with at least 5 CpG site overlaps with the reference cell-type atlas were eligible to be mapped to a particular cell type, the resulting cell type-specific long-read methylation profiles contained a small number of intersecting CpG sites (Figure 3E), necessitating a reference-based aging model that uses different sets of CpGs for each sample. Applying LongReadAge to these cell type-specific methylation profiles demonstrated a robust ability to track aging across both myeloid cells and lymphocytes, with mean errors below 6 years for both cell types (Figure 3F). Thus, long-read sequencing data has the potential to resolve epigenetic ages within a bulk sample at cell-type resolution. Comparing the epigenetic age estimates between myeloid and lymphocyte cells across the same individuals, we found there was no significant difference between the two (Figure 3G).

### LongReadAge captures aging of circulating cell-free DNA

Circulating cell-free DNA (cfDNA) consists of DNA fragments outside of cells that circulate in plasma and other biofluids, often as a result of cell death. Molecular alterations in cfDNA are observed with aging and may serve as valuable biomarkers of the aging process^26,27^. We applied LongReadAge to track epigenetic aging in long-read plasma cfDNA data derived from two cohorts: one from the Chinese University of Hong Kong (CUHK)^17,18^ and another from the Mass General Brigham (MGB) Biobank, which was newly generated in this study (Figure 4A). To generate this data, we obtained plasma samples from the MGB Biobank, extracted cfDNA, and profiled long-read methylation using PacBio Single Molecule Real-Time (SMRT) sequencing. Across both cohorts, LongReadAge demonstrated a modest ability to track aging (Figure 4B).

**Figure 4:**
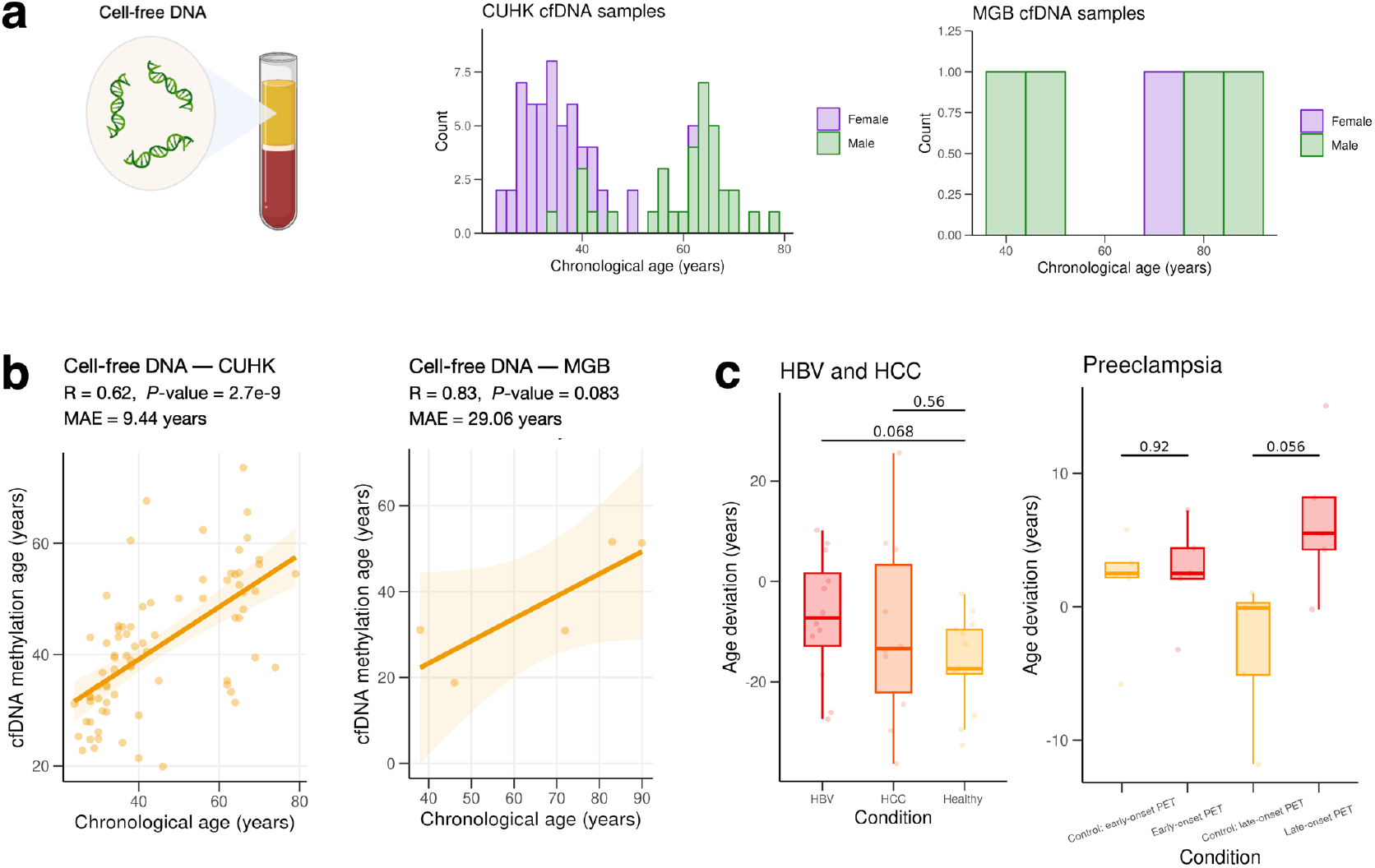
LongReadAge captures aging of circulating cell-free DNA. **(a)** Cell-free DNA derived from plasma was profiled in two cohorts: one from the Chinese University of Hong Kong (CUHK) consisting of 84 samples (healthy + pregnancy + disease) and one from the Mass General Brigham (MGB) Biobank (5 samples). **(b)** Application of LongReadAge to CUHK and MGB cohort. Samples with fewer than 300 CpG sites intersecting the reference CpGs were omitted. **(c)** Comparison of age deviation between 1) hepatitis B virus (HBV) carriers, hepatocellular carcinoma (HCC) patients, and healthy controls and 2) early-onset and late-onset preeclampsia (PET) vs respective controls.

The CUHK cfDNA cohort included patients with a broad spectrum of diseases and health conditions, including hepatitis B virus (HBV) carriers, hepatocellular carcinoma (HCC) patients, healthy controls, and cases of early-and late-onset preeclampsia (PET) with respective controls. When comparing the cfDNA age deviation (predicted age minus chronological age) between HCC subjects and controls, as well as between HBV subjects and controls, we observed a marginally significant increased age deviation in the HBV group compared to the healthy group and no difference between HCC patients and healthy controls (Figure 4C). In the comparison between PET subjects and controls, no differences were found in early-onset PET cases, while a marginally significant increase in cfDNA age deviation was observed in late-onset PET cases (Figure 4C).

## Discussion

Understanding cell-type biology has been indispensable in characterizing various age-related diseases, from dopamine-neuron-specific deterioration in Parkinson’s disease^28^ to beta-cell-specific dysfunction in type 2 diabetes^29^. A comprehensive understanding of aging will require similarly granular elucidation of how individual cell types deteriorate over time, and how cell-specific changes relate to aging at tissue, organ, and organismal levels. Presently, however, options for profiling cell type-specific features of aging are limited, and our understanding of aging trajectories between different cell types remains incomplete.

In this study, we present a novel framework, LongReadAge, that predicts the epigenetic age of sparsely intersecting long-read methylation profiles. LongReadAge shows robust performance in capturing aging of peripheral leukocytes, and its reference-based architecture enables accurate age profiling despite limited numbers of long-read samples and limited shared CpGs across samples.

From bulk leukocyte long-read data, we generated immune-cell-specific methylation profiles that demonstrated a strong concordance with reference cell types. Extending LongReadAge to these methylation profiles revealed robust performance in profiling cell type-specific aging despite vanishingly low numbers of shared CpG sites between samples. We further demonstrated the ability of LongReadAge to capture epigenetic aging of plasma cfDNA. While the performance is more modest here, the observed marginally significant increase in cfDNA age deviation in disease states, including hepatitis and preeclampsia, suggests potential applications of our model to evaluating aging in the context of disease.

While our study focuses on peripheral blood, the versatility of our approach enables its applications to additional tissues and disease states. It could be used, for instance, to study neuron-specific aging in long-read brain data, or endothelial cell-specific aging in long-read heart data. Another promising application of this approach is tracking organ-specific aging through plasma cfDNA. As cells die throughout the body, their DNA is released into the bloodstream as circulating cfDNA. Thus, cfDNA comprises DNA from various organs throughout the body, a feature leveraged by recently developed liquid biopsies, which have shown promise in diagnosing early-stage solid-tissue cancers based on epigenetic signals from plasma cfDNA^30–32^. Extending this concept, our approach could be applied to infer the epigenetic age of different solid-tissue organs, tissues, cell types, and perhaps individual cells, from a single blood sample. As long-read sequencing throughput continues to improve and comprehensive reference datasets become more widely available, the prospect of accurately determining the epigenetic age of multiple cell types, including those from solid organs, from a single blood draw is becoming increasingly feasible. This approach could revolutionize our ability to monitor aging at organ-, tissue-, and cell-specific resolution.

Looking ahead, several key steps could enhance this approach. First, developing supervised machine learning models to predict the origin of DNA fragments could improve the robustness of cell-type assignment and may enable more fragments to be mapped to their origin cell type^19^. Second, employing whole-genome reference aging datasets would greatly expand the capabilities of our approach. Currently, our EPIC array-based aging reference covers only 3% of the genome, leaving the vast majority of CpG sites measured in long-read sequencing ineligible for age prediction. Access to whole-genome reference aging datasets would allow us to profile a broader range of cell types and may even enable age prediction at the level of individual DNA fragments. While whole-genome cell type-specific methylation atlases, such as those developed recently^23^, do exist, their current sample sizes and age ranges are inadequate for use in our model as an aging reference dataset. However, with the rapidly declining costs of DNA sequencing, generating comprehensive, whole-genome, cell type-specific references will likely become increasingly feasible in the future.

Several limitations to this study should be noted. First, our reference aging methylation dataset was derived from whole blood samples. To better capture cell type-specific aging, the reference aging dataset would ideally be cell-specific as well (e.g., a myeloid cell aging reference for myeloid cell long-read methylation profiles). While such references do not exist presently, future incorporation could improve the ability of our approach to capture cell type-specific features of aging. Second, our model currently measures only linear changes corresponding to the passage of chronological time. While these changes serve as a reasonable first approximation of aging, future models could be constructed to better capture functional fitness of individual cell types or cell-specific deleterious changes. Third, the cohort in which our cell type-specific model is validated is limited in demographic diversity, consisting entirely of pregnant women around the age of 20 to 40 years. Validating this framework in more diverse cohorts will be essential in the future.

In sum, our study introduces a novel approach for profiling epigenetic aging at cell-type resolution using long-read methylation sequencing. This method opens new avenues for investigating cell type-specific aging across various cell types and tissues. As long-read sequencing technologies continue to advance and reference datasets improve, we anticipate that this approach will become an increasingly valuable tool in aging research and biomarker development. Future applications may provide valuable insights into cell type-specific aging in both health and disease and facilitate the development of increasingly biologically precise aging biomarkers.

## Methods

### Ethics

This study was approved by the Mass General Brigham Institutional Review Board under Human Protocol #2021P003059. Details of ethics approval for previously generated datasets used in this study are found in each dataset’s respective publication: ^17,18,22,23,30^.

### Long-read datasets

Our study utilized long-read datasets from the Chinese University of Hong Kong, which are detailed in publications by Choy et al., 2022^17^, Yu et al., 2023^30^, and Yu et al., 2021^18^. Across these datasets, we used 16 long-read buffy coat samples and 84 cfDNA samples. For long-read data, DNA was extracted from buffy coat and plasma using Qiagen’s QIAamp DNA Blood Mini Kit and QIAamp Circulating Nucleic Acid Kit, respectively. Long-read sequencing was then conducted on PacBio’s Sequel II platform using the SMRT Cell 8M and the SMRTbell Express Template Prep Kit 2.0.

### Microarray datasets

For our reference aging dataset, we used a blood methylation dataset available on GEO under accession GSE152026, as described in Hannon et al., 2021^22^. Briefly, DNA was extracted from whole blood, bisulfite-converted using the EZ-96 DNA Methylation-Gold Kit, and methylation was profiled on the Illumina EPIC microarray platform. We filtered this dataset to include only control subjects, resulting in 519 samples.

### Whole-genome bisulfite sequencing datasets

As our cell type-specific methylation atlas, we employed whole-genome methylation sequencing data available on GEO under accession GSE186458 and described in more depth in Loyfer et al., 2023^23^. Specifically, we utilized 36 methylation profiles measured in different blood cell types.

### MGB long-read cfDNA cohort

Our study generated a new dataset profiling cfDNA methylation using PacBio long-read technology. Plasma samples and health information were obtained from the Mass General Brigham (MGB) Biobank, a biorepository of consented patient samples at MGB, from healthy subjects spanning a broad age range. DNA was extracted from 7-8 ml of plasma using Qiagen’s Circulating Nucleic Acid Kit, and cfDNA concentration and genomic DNA contamination were assessed using the Agilent TapeStation 4150 automated electrophoresis platform and Agilent Cell-free DNA ScreenTape. DNA was filtered to enrich for longer fragments using Zymo’s Select-a-Size MagBead Kit.

For long-read sequencing, five plasma DNA samples (ranging from 34-75 ng of DNA) were sequenced on PacBio’s Sequel II platform using Single Molecule Real-Time Sequencing. Samples were treated with RNase to eliminate RNA contamination and concentrated using 3x AMPure Beads. Then, libraries were prepared with PacBio’s SMRTbell Template Prep Kit 3.0, samples were barcoded and pooled with two or three samples per pool, and polymerase was bound to templates using the Sequel II Binding Kit 3.2. Finally, samples were multiplexed onto two SMRT Cells (8M) and sequenced on the Sequel II with 30-hour movie times.

### Long-read processing

All long-read data were processed using PacBio’s SMRT Tools software (v12.0). Raw reads (subreads) were converted to circular consensus sequence (CCS) reads using the *ccs* tool, with the hifi-kinetics parameter set to output fluorescence kinetics signals, which are used to infer CpG methylation. For the MGB data, samples were demultiplexed using *lima* along with their respective barcodes. CpG methylation information was extracted from the kinetic signals using the *primrose* tool, and the CCS reads were aligned to the hg19 reference genome using *pbmm2 align* under the CCS preset. The CCS reads were then converted to BED files describing CpG methylation across the genome using *pb-CpG-tools* (v2.3.1) with a pileup mode of count and a minimum coverage of 1.

### LongReadAge model development

LongReadAge is based on the scAge framework. Like scAge, we employed a reference methylation dataset, referred to here as the reference aging dataset, and built linear regression models to predict CpG methylation as a function of age across every CpG site in the reference aging dataset. Our reference was constructed using blood EPIC microarray data. The input to our model is a long-read methylation profile, consisting of a series of CpGs and their methylation statuses, which range from 0 to 1 and represent the proportion of CpGs with a methyl group. These CpGs were intersected with CpGs in the reference EPIC dataset and filtered to include only the top age-correlating CpG sites.

To select the top age-correlating CpG sites, we identified the top 1% of CpGs with the highest absolute Pearson R correlation coefficient with age, unless otherwise noted. For cell-specific methylation profiles, we observed better performance when using a fixed number of top CpG sites rather than a percentile cutoff. The fixed number of CpG sites was determined by finding the mean number of CpGs used at a 1% percentile cutoff. For the cfDNA samples, several had fewer than 300 intersecting, top-correlating CpG sites at a 1% percentile cutoff. These samples were removed from further analysis.

Using the top-correlating CpG sites, we predicted methylation as a function of age across ages ranging from 0 to 100 years in increments of 0.1 years. For each predicted methylation pattern, we determined how well it matched the observed methylation in the long-read methylation profile by computing a probability score using the following formula, where *n* is the number of top age-correlating CpG sites in the intersected methylation profile, *k* is a particular CpG site index, *p* is the predicted methylation vector, and *m* is the observed methylation vector.

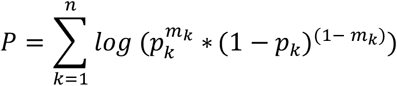

The age that best predicted the observed methylation pattern, yielding the highest probability score, was designated as the sample’s epigenetic age. Further details of this algorithm are described in Trapp et al., 2021^13^.

### Generating cell type-specific methylation profiles

To generate cell type-specific methylation profiles from bulk long-read methylation data, we developed a three-step process. First, differentially methylated regions were identified between cell types of interest. Second, long-read fragments were traced to their cell type of origin. Third, methylation profiles were generated from all reads predicted to originate from a given cell type. This process was based on methods developed by Choy et al., 2022^17^ and Yu et al., 2021^18^, with several refinements.

#### Identifying differentially methylated regions

We first identified differentially methylated regions between cell types of interest by leveraging a cell type-specific reference methylation atlas published in Loyfer et al., 2023^23^ and available at GSE186458. Specifically, we used the 36 samples measured in blood encompassing all major leukocyte cell types.

We categorized each sample into its respective target cell type (myeloid or lymphocyte) (as described in Table S1), then found genomic regions whose methylation patterns differentiated these cell types by applying the software suite *wgbstools* ^24^. This process involved two steps: first, identifying genomic segments (or blocks) of similarly methylated regions based on input *beta* files with *segment*, then finding differentially methylated blocks using *find_markers*, which applied several criteria to identify robust differentially methylated regions, including cutoffs for mean methylation, quantiles, t-test p-value, and coverage. The exact parameters used for these steps are listed in Table S2.

Once these markers were determined, they were mapped from genomic blocks to individual CpGs to enable CpG-specific comparison with long-read fragments, yielding a final table with mean methylation of each cell type across each differentially methylated CpG site, referred to here as the reference cell-type atlas. Of note, a reference cell-type atlas is generated for each cell type of interest.

#### Tracing fragments to their cell type of origin

To identify long-read fragments’ origin cell type, we sought to determine how well a given fragment matched the methylation of a particular cell type in the reference cell-type atlas. To achieve this, we first filtered the fragments to include only autosomal fragments with a minimum mapping quality of 20.

We then identified the long-read fragments with at least 5 CpGs that overlapped with the reference atlas for each cell type. We chose the value 5 as the minimum overlap based on a computational analysis conducted by Yu et al., 2021^18^ that found 5 sites as the minimum for area under the receiver operating characteristic (AUROC) values exceeding 82% in performing tissue-of-origin determination. From these overlapping CpGs, we computed a likelihood score describing how well the long-read fragment matched each cell type in the reference cell-type atlas. The likelihood score was calculated using the following formula where *n* is the number of overlapping CpGs, *k* is a particular CpG site index, *r* is the long-read methylation profile, and *c* is the cell-type methylation profile in the reference cell-type atlas.

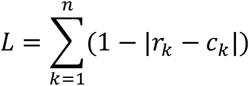

Fragments with a likelihood score for one cell type at least (1 / 0.95) times greater than all other cell types were considered to originate from that cell type.

#### Generating cell type-specific methylation profiles

Finally, we filtered the original CCS long-read files to include only reads predicted to originate from a given cell type using Picard’s *FilterSamReads* and generated methylation profiles from the resulting reads using *pb-CpG-tools* (v2.3.1) with a pileup-mode of *count* and a minimum coverage of 1.

### Quantification and statistical analysis

All statistical analyses were performed in R (version 4.0.2) and Python (version 3.12). Correlations were computed using the Pearson method, and a Wilcoxon signed-rank test was used to determine the statistical significance of differences between groups. Sample sizes were based on the number of replicates in each dataset, and no power analyses were conducted. A *P*-value threshold of 0.05 was used for determining significance.

## Acknowledgments

We thank the Yale Center for Genome Analysis (YCGA) and Keck Microarray Shared Resource (KMSR) at Yale University for providing the necessary PacBio sequencing services, which is funded in part by the National Institutes of Health instrument grant 1S10OD028669-01.

This study makes use of data generated by The Chinese University of Hong Kong (CUHK) Circulating Nucleic Acids Research Group, as reported by Choy et al in Clin Chem (DOI: 10.1093/clinchem/hvac086), Yu et al in Clin Chem (DOI: 10.1093/clinchem/hvac180), Yu et al in Proc Natl Acad Sci USA (DOI: 10.1073/pnas.2114937118).

## Availability of data and materials

The CUHK datasets are not publicly available; however, they can be requested by contacting the CUHK Circulating Nucleic Acids Research Group. Data from the newly generated MGB cfDNA dataset will be made available upon publication.

## Code availability

Code will be made publicly available upon publication.

## Authors’ contributions

A.E. extracted and sequenced cfDNA in the MGB cohort, obtained access to and processed CUHK datasets, developed the LongReadAge model, performed all analyses, and drafted the manuscript. M.M. guided computational analyses and contributed to manuscript preparation. J.P. contributed to sample procurement and assisted with manuscript writing. V.N.G. conceived the study, oversaw research, and wrote and revised the manuscript.

